# CRAST Leads to Homologous-ncRNA Search in Genomic Scale

**DOI:** 10.1101/127738

**Authors:** Masaki Tagashira

## Abstract

**Motivation:** Non-coding RNAs (ncRNAs) play important roles in various biological processes. In past, homologous-ncRNA search in genomic scale (e.g., search all house mouse ncRNAs for several human ones) is difficult since explicit consideration of secondary structure in alignment leads to impractical complexity on both of time and space.

**Results:** In this study, building the program *CRAST* (Context RNA Alignment Search Tool, available at “https://github.com/heartsh/crast” including the used validation/test set), we developed the CRAST algorithm, a *“seed-and-extend”* alignment one based on adaptive seed and RNA secondary structure context (motif probabilities) as in Fig. 1. The algorithm is *O*(*n*: *a sum of lengths of target sequences*) on time through help of adaptive seed, implicitly considering both of sequence and secondary structure; it provides computation time comparable with other BLAST-like tools, significantly reduced from any variant of the Sankoff algorithm for alignment with the explicit consideration. It detects homologs as many as other BLAST-like tools and the lowest number of non-homologous ncRNAs.

## 1 INTRODUCTION

NcRNAs are involved in diverse biological processes such as rRNA modification [1] and chromatin modification [2]. RNAs including ncRNAs prefer to form into 3D structures rather than DNAs due to contribution of 2’ hydroxyl group to hydrogen bond and function based on it. Hence only sequence identity doesn’ t lead to ncRNA function. [3]

**Figure 1:**
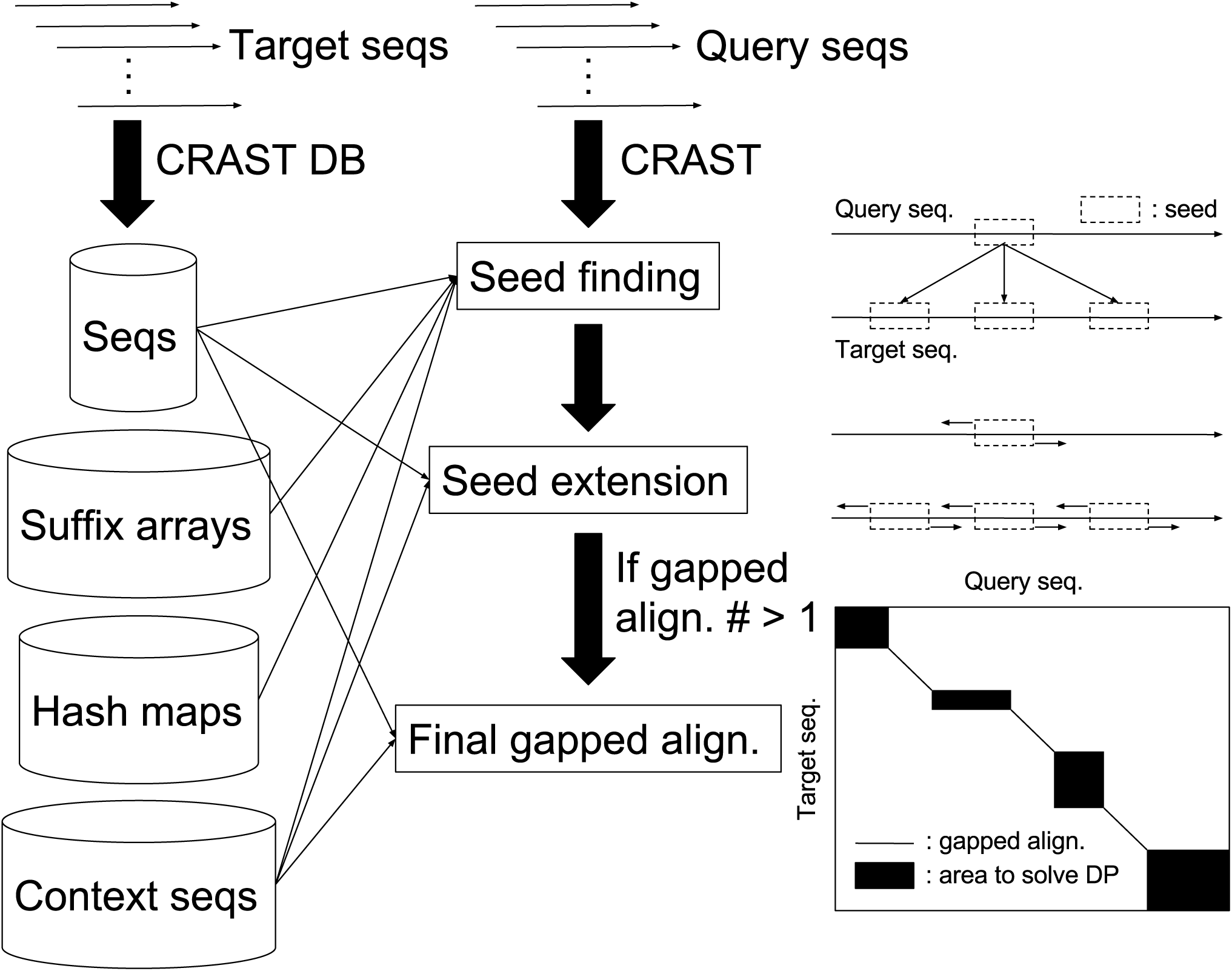
overview of CRAST

### 1.1 General genomic homolog-search

In genomic DNA/protein-homolog-search, tools based on “seed-and-extend” strategies such as BLAST [4] and LAST [5] are mostly used. In summary, these tools find *seeds* based on sequence identities of partial regions of 2 compared sequences, and extend the seeds from both sides of them til their scores based on sequence identities drop to some extent. They reduce the time complexity *O*(*mn*) from the optimal search by the Smith-Waterman algorithm [6] with the suboptimal search.

### 1.2 Genomic ncRNA-homolog-search

On the other hand, secondary structure identity must be taken into account as well as sequence identity in genomic ncRNA homolog search from the aforementioned matter (especially for ncRNA with low sequence identity such as lncRNA). [7] Foldalign, an implementation of the Sankoff algorithm [8] finds ncRNA pairwise alignments by simultaneously folding and aligning sequences with pruning of dynamical programming matrix. [9] It is really time-consuming and not applicable to genomic data (e.g., all house mouse ncRNAs to search for several human ones) although its pruning discards any subalignment that doesn’ t have a score above a length-dependent threshold. The banded Sankoff algorithm, another variant of the Sankoff algorithm reduces the time/space complexity of the Sankoff algorithm *O*(*n*^6^)/*O*(*n*^4^) to *O*(*n*^4^)/*O*(*n*^3^), which again results in unapplicability to genomic data. [10] After all, explicitly considering RNA secondary structure and sequence results in the huge complexities on both time and space.

We could implicitly and efficiently consider RNA secondary structure using CapR, a linear time/space complexity (*O*(*nw*^2^)/*O*(*nw*), but *w* could be considered as a constant) algorithm to estimate an RNA secondary structure context (motif probabilities) of each base in any RNA [11]. This probabilistic encoding of RNA secondary structure seems to enable to align ncRNA sequences in the same fashion as alignment for DNA/protein sequence.

So we built CRAST, a BLAST-like genomic ncRNA alignment search tool. We demonstrated this tool enables to align ncRNAs with consideration of secondary structure and sequence in a time complexity *O*(*n*) where *n* is a sum of lengths of target sequences through helps of adaptive seed adopted in LAST [5], suffix array [12], and CapR.

## 2 METHODS

We implemented CRAST in Rust, a systems programming language [13]. It supports both of safe parallelism (multithreading without data race) and safe zero-cost abstraction (e.g., runtime without garbage-collection and guarantee of memory safety when compiling), which results in both of more safety and computation efficiency almost the same as/more than C/C++. We implemented the bottlenecks (e.g., the calculation of the context sequences and alignment search) in a multi-threaded fashion.

### 2.1 NcRNA seed finding

“Seed-and-extend” using fixed-length seed such as one in BLAST leads to a quadratic number of seeds with target sequence length due to a non-uniform (oligo-)nucleotide composition of any real sequence. [5] Adaptive seed to find matches that occur at most *f* times in any target sequence guarantees a linear number of seeds and linear time complexity with a target sequence length. [5] We adopted this seed for extracting matches without need of repeat-masking which could hide potentially significant parts.

To find adaptive seeds, we initially build suffix arrays of target ncRNAs in a time/space complexity *O*(*n*). Then we generate hash maps of short substrings of target sequences to corresponding index ranges in the suffix arrays for access in a time complexity *O*(1). We store the suffix arrays, hash maps, target sequences, and context sequences of target sequences in compressed files (of the “bzip2” format) as a database.

We find the shortest seeds for starting position pairs of any query sequence and target one with the hash maps for short matches; with binary-search using search ranges found in previous searches for long ones, as in [5]. These 2 techniques and search strategy reduce the time complexity *O*(*m* log *n, m*: *a length of a match*) of the binary-search for the substring search using suffix array into one less than *O*(log *n*). Steps til here are the same as in LAST, an adaptive seed implementation [5] except for the context sequences.

In CRAST, the seeds having similar context sequences are filtered in from the found seeds, using a threshold of an expected number of the seeds with more/equally similar context sequences as in Fig. 2.

**Figure 2:**
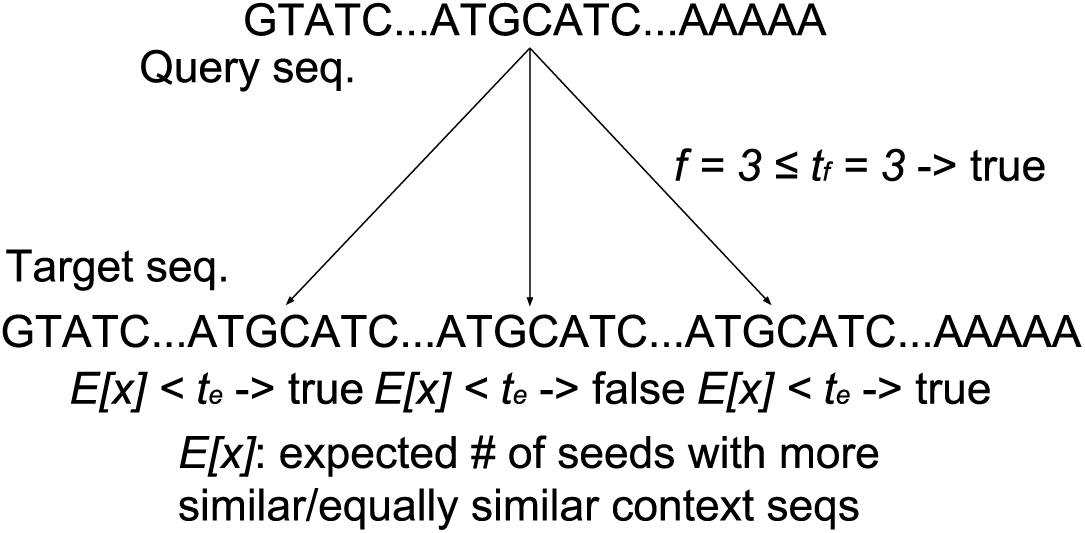
seed finding of CRAST The seeds only satisfying both of the terms for sequence/context sequence match (the left/right one) are filtered in.

We score any base pair having the Jensen-Shannon distance [14] of a pair of contexts as in Fig. 3 less than 0 < *p* < 0.5 as a match with a score +1; other than that as a mismatch with a score -1. The distance is a distance metric version of the Jensen-Shannon divergence [15] (not distance, often called the JSD) which is a symmetric finite measure of similarity between 2 probability distributions. The JSD doesn’ t satisfy triangle inequality required for any distance metric while the distance does. We considered a binomial distribution as a model of a probabilistic distribution of the series of the matches and mismatches such as a series of coin tosses: *B*(*n,* 0 < *p <* 0.5).

**Figure 3:**
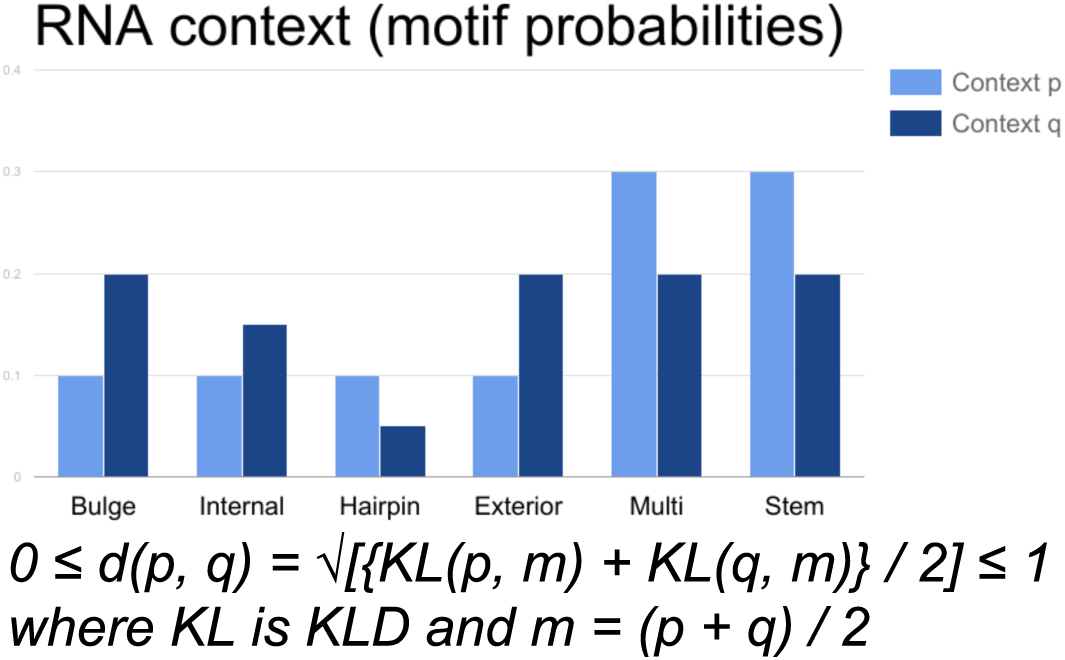
Jensen-Shannon distance of RNA context pair Each term for the context is an RNA secondary structure motif.

### 2.2 Seed extension and scoring system

We score any base pair using both of match/mismatch of base/context: *s* = *rs*_*b*_ + (1 – *r*)*s*_*c*_ where *s* is a fusion score, 0 ≤ *r* ≤ 1 is a contribution ratio of base to the fusion score, *s*_*b*_ is a score of any base pair (+1/-1), and *s*_*c*_ is a score of any context pair (+1/-1) in the CRAST scoring system.

We extend the seeds first without any gap using the X-drop algorithm [16]; then with gaps using the same algorithm. The algorithm greedily extends the seeds/ungapped alignments til their scores drop from maximum observed values *m* to less than/equal to *m − x* where *x* is an aforehand determined value as in Fig. 4. [16]

**Figure 4:**
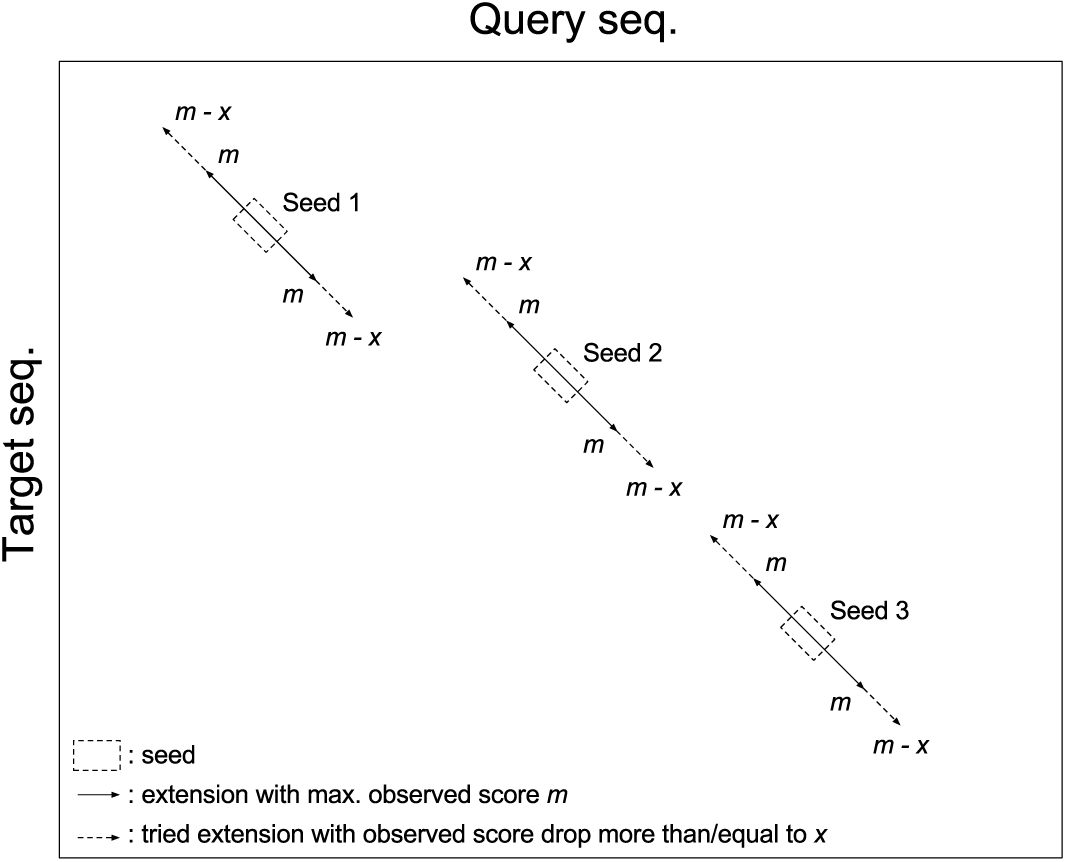
X-drop algorithm

Finally, we generate single gapped alignments derived from all the gapped ones, using DP matrices constrained by the gapped ones as in [17] and Fig. 5 when a number of the gapped ones is more than 1 for 1 strand. To constrain the matrices, we greedily merge the ungapped ones diagonally overlapped in the matrices; greedily remove one with a lower score of the ungapped ones non-diagonally (in parallel) overlapped. We greedily remove one with a lower score of the overlapped gapped ones for the same purpose. After each of the ungapped/gapped one, we discard some of the alignments based on an expected number of alignments with more/equally similar base/context sequences.

**Figure 5:**
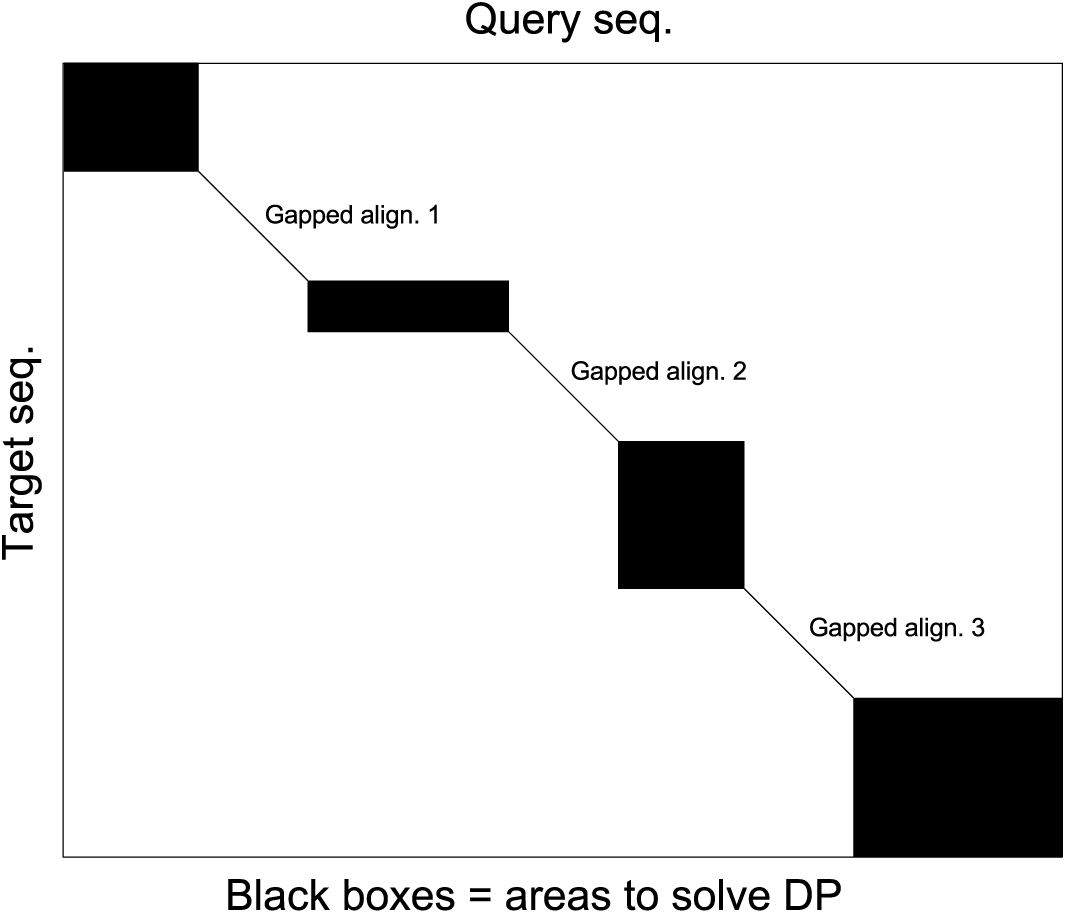
constrained DP matrix of CRAST All the area of the DP matrix to solve the DP are reduced from *mn*.

More specifically, we independently consider a binomial distribution of the series of the base/context matches and mismatches, which results in simple calculations of the E-values. The gapped one disables to model the distributions due to uncertainty of gap; we consider the gaps as given, which results in the same calculations of the E-values as the ungapped one.

## 3 RESULTS

### 3.1 Parameter-tuning

We set default CRAST parameters as in Table 3 to best perform with target/query sequences for validation. We used all 18,185 *Mus musculus* ncRNAs (derived from Ensembl [18]) as the target sequences; 34 *Homo sapiens* lncRNAs known as homologs to *M. musculus* corresponding ones including HOTAIR [19] and XIST [20] (derived from LncRNAdb [21]), as the query sequences. We fixed the parameters not referred to in each result.

### 3.2 Relation between match probability of RNA context and homolog detectability

Fig. 6 shows the higher the match probability of RNA context *p* becomes, the more the FPs are obtained while the number of the TPs hits the ceiling at *p* = 0.25.

**Figure 6:**
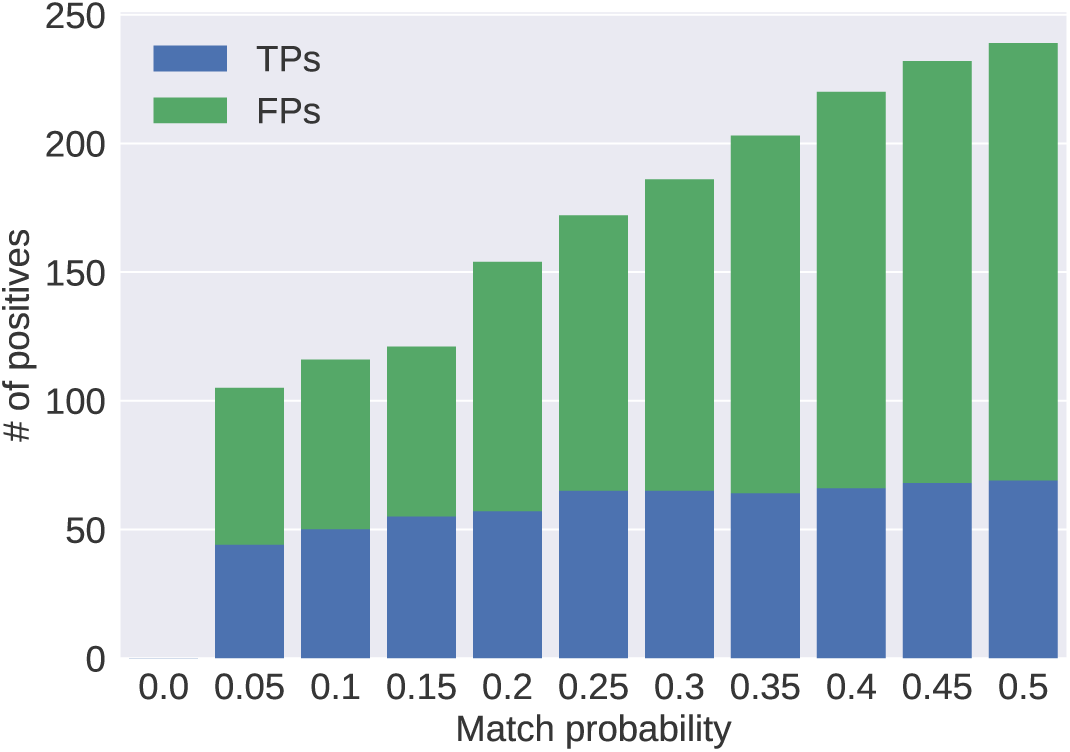
relation between match probability of RNA context and homolog detectability We used the same target/query sequences for test as the target/query sequences for validation. The “TP” is map of any *Homo sapiens* one to any corresponding *Mus musculus* one; the “FP” is any *Homo sapiens* one to any of the others.

### 3.3 Relation between seed E-value filtering and homolog detectability

Fig. 7 shows the number of the FPs hits the ceiling at *t*_*e*_ = 1.0 · 10^4^ while the one of the TPs (homologs) hits the ceiling at *t*_*e*_ = 1.0 · 10^3^ where *t*_*e*_ is any threshold of an expected number of the seeds with more/equally similar context sequences. From that, we could think *t*_*e*_ as a parameter to discard the inappropriate seeds before the following extensions. It also implies only the seeds with highly similar context sequences would lead to incomplete capture of homologs.

**Figure 7:**
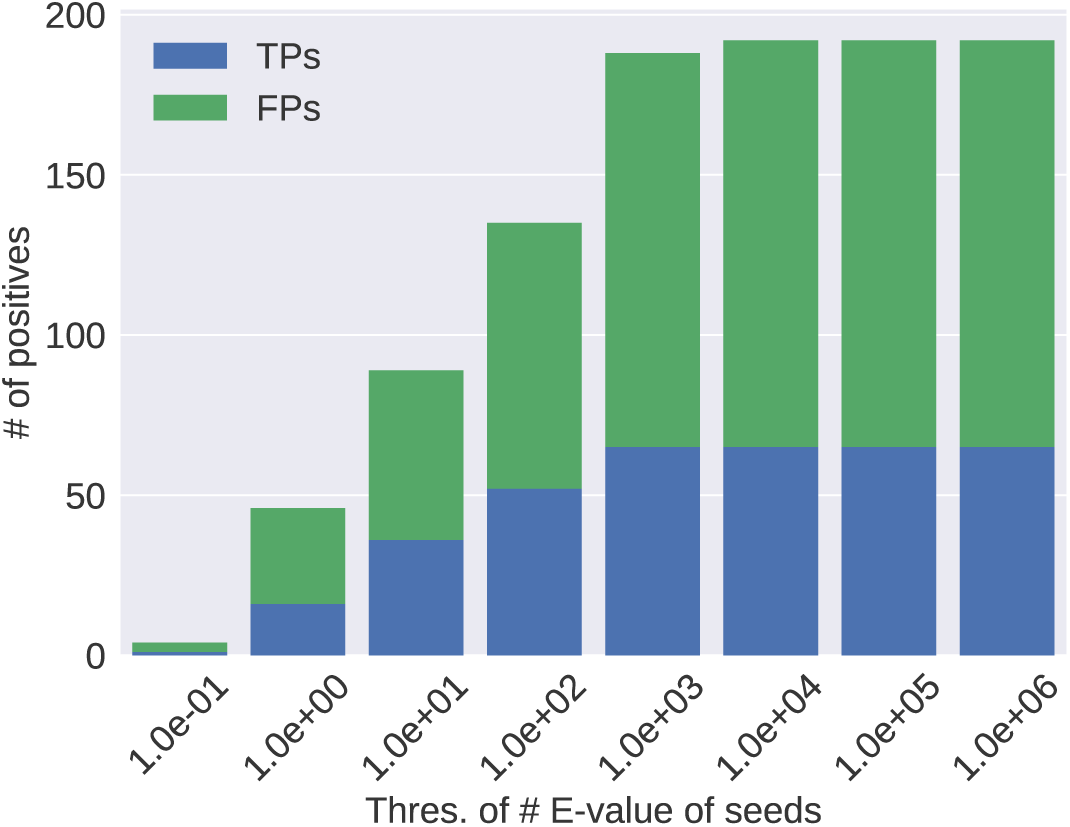
relation between seed E-value filtering and homolog detectability We used the same target/query ncRNAs as Fig. 6.

### 3.4 Relation between contribution ratio of base to score *r* and homolog detectability

Fig. 8 represents the higher the contribution ratio of base to the fusion score *r* becomes, the more the FPs are obtained while the number of the TPs hits the ceiling at *r* = 0.7. From that, we could think *r* as a parameter to control a number of the FPs keeping the TPs as many as possible. It also implies consideration of only secondary structure would lead to incomplete capture of homologs.

**Figure 8:**
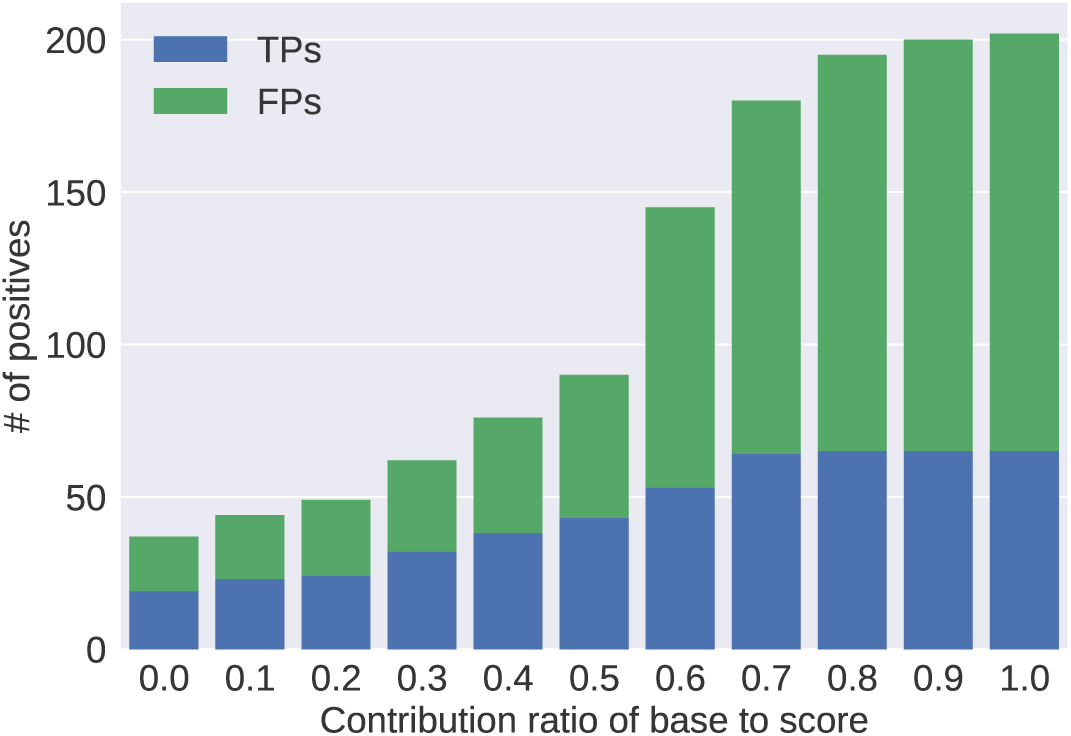
relation between contribution ratio of base to score *r* and homolog detectability We used the same target/query ncRNAs as Fig. 6.

### 3.5 Relation between alignment E-value filtering

Fig. 9 shows the number pair of the TPs and FPs hits the ceiling around *t*_*e*1_ = 1.0 · 10^−3^ and *t*_*e*2_ = 1.0 · 10^3^ where *t*_*e*1_/*t*_*e*2_ is any threshold of an expected number of the alignments with more/equally similar sequences/context sequences. From that, it is confirmed that each of the Evalue thresholds could play a role as a parameter to discard the inappropriate alignments before the following alignment step/report. It also implies only the alignments with highly similar context sequences would lead to incomplete capture of homologs as well as the seeds.

**Figure 9:**
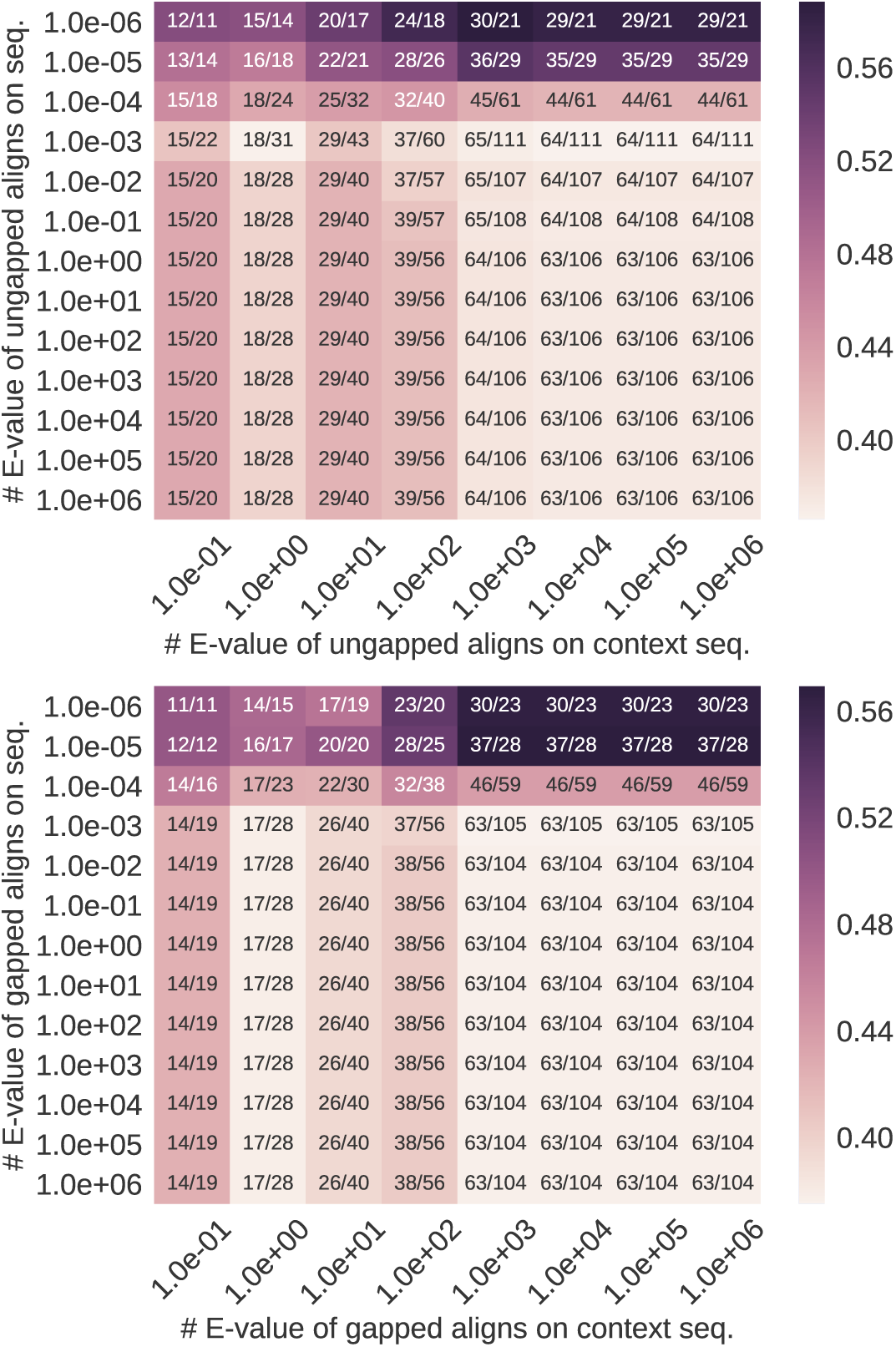
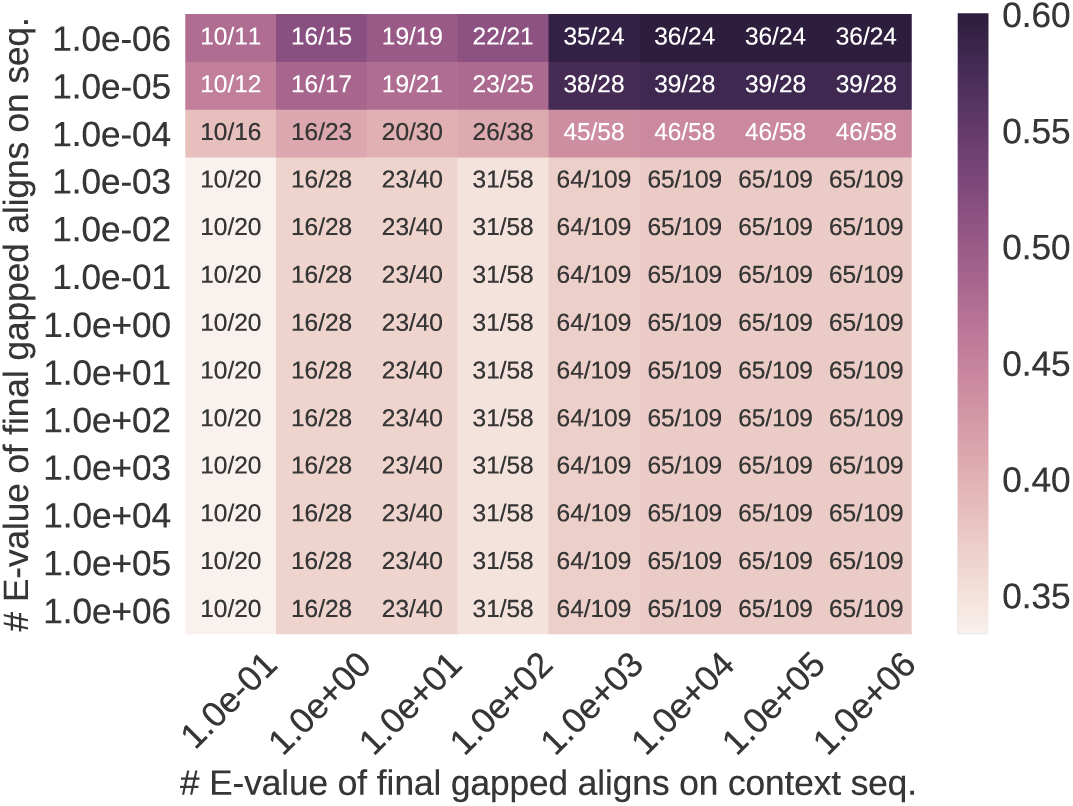
relation between alignment E-value filtering and homolog detectability We used the same target/query ncRNAs as Fig. 6. From the top, the relation for: ungapped one/gapped one/final gapped one. A color of each cell indicates precision *p* = *T P/*(*T P* + *F P*); a number of each one does TP/FP number.

### 3.6 Performance comparison with BLAST-like tools

We compared CRAST, LAST (ver. 719), and BLASTN (BLAST for nucleotide, ver. 2.6.0+) for effect on ncRNA homolog detection of fixed-length/adaptive seed/seed-extension based on only substitution matrix/both of substitution matrix and RNA secondary structure context as in List. 1. We also measured running times of these programs on both database/alignment step. (In practice, database step is run only 1 time for any set of target sequences.)

Table 1 shows CRAST detects the homologs as many as the compared tools and the lowest number of the FPs. It also shows the database step of CRAST is significantly timeconsuming due to domination by the CapR algorithm; the alignment step is relatively slow in spite of a lower number of the seeds.

**Table 1:**
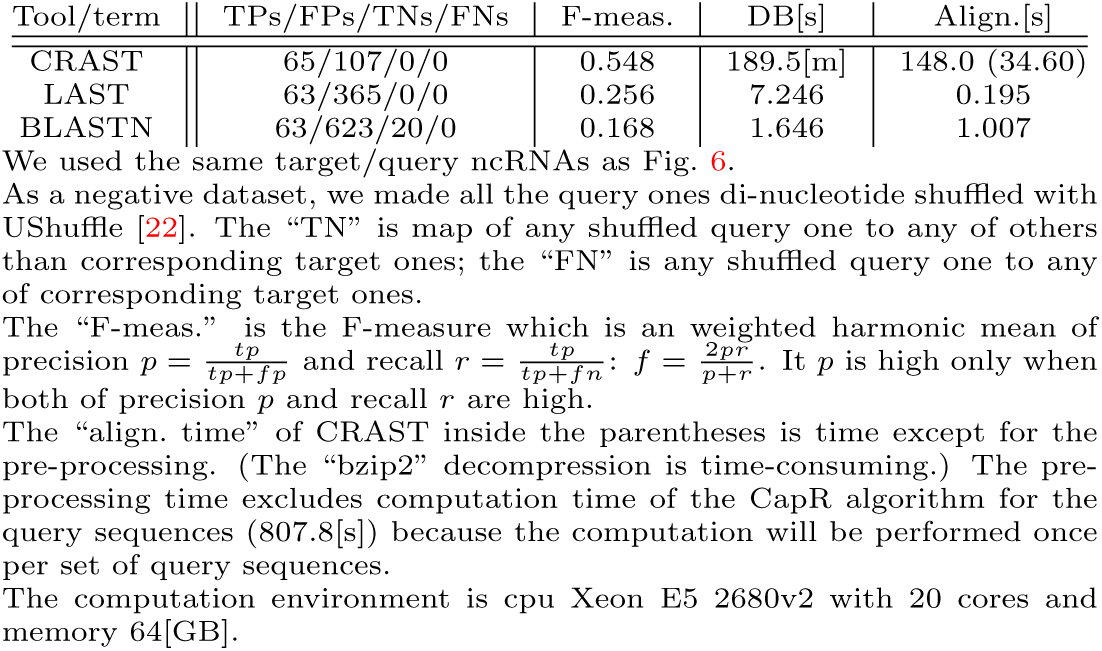
performance comparison among CRAST, LAST [5], BLASTN [4] We used the same target/query ncRNAs as Fig. 6.

3.6 Performance comparison with BLAST-like tools
| Tool/term | TPs/FPs/TNs/FNs | F-meas. | DB[s] | Align.[s] |
| --- | --- | --- | --- | --- |
| CRAFT | 65/107/0/0 | 0.548 | 189.5[m] | 148.0 (34.60) |
| LAST | 63/365/0/0 | 0.256 | 7.246 | 0.195 |
| BLASTN | 63/623/20/0 | 0.168 | 1.646 | 1.007 |

There are 2 reasons why the alignment step is relatively slow: missing pre-computing of seed candidates (possible adaptive seeds) in the database step and the compute-intensive of the Jensen-Shannon distance. First, we could reduce computation time of the alignment step by precomputing the candidates in a certain database within a given range of seed frequencies (e.g., the frequencies around default one), which would lead to search of the candidates in query sequences, not target ones. But this pre-computation doesn’ t repesent the real performance of the CRAST algorithm because the speed-up is established only in the case when seed frequencies are within the range. Second, the Jensen-Shannon distance is computationally intensive compared with the *L*_*p*_ one such as the Euclidean one (*p* = 2) since it involves the intensive computation of logarithm compared with basic arithmetic operations. Considering these 2 reasons, the computation time of CRAST is reasonable.

Fig. 10 shows the ROC curves of CRAST and the other BLAST-like tools using their alignment expectations as the thresholds. (CRAST emits both of the expectations based on sequence/context sequence, however, we used only the ones based on sequence for the ROC curve.) Its shows the prediction performance of CRAST is superior to the other tools.

**Figure 10:**
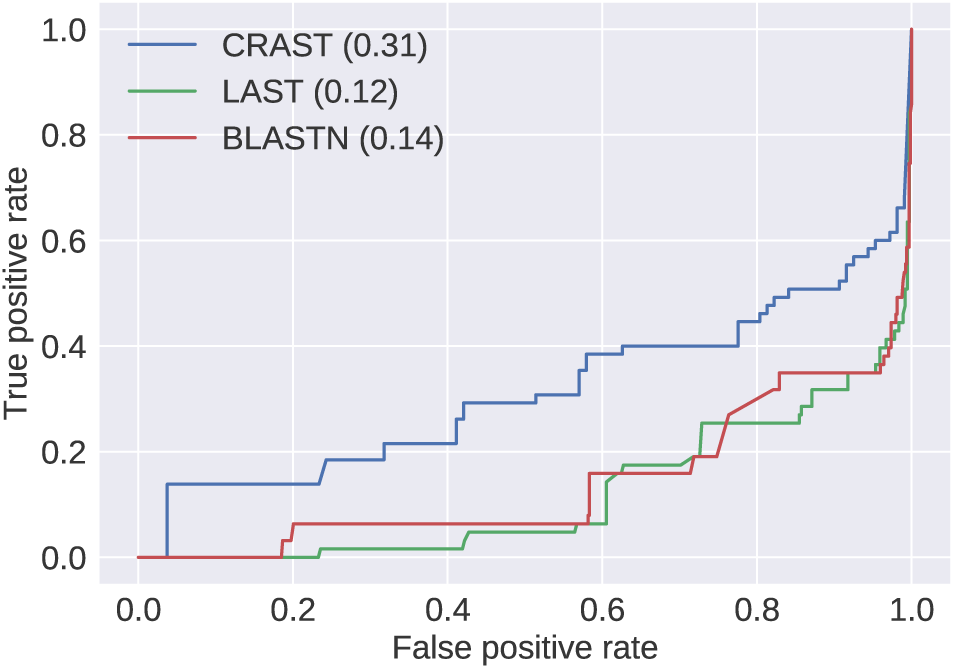
ROC curves of CRAST, LAST, and BLASTN The used dataset and setting of the binary classification are the same as Table 1. The curves are derived using their alignment expectations as the thresholds. (CRAST emits both of the expectations based on sequence/context sequence, however, we used only the ones based on sequence for the curve.) Values inside the parentheses are areas under the curves. (The larger the area becomes, the better the prediction performance gets.)

### 3.7 Performance comparison with Foldalign

We compared CRAST and Foldalign (ver. 2.5) for effect on ncRNA homolog detection of alignment with context sequence/simultaneous folding as in List. 2.

Table 2 represents there are numerous incorrect maps of the target ones to others than the corresponding query ones for Foldalign (32 + 1, 923 – 219 = 1, 768) compared with CRAST (65 + 156 – 219 = 2); a half of correct maps of the target ones to the corresponding ones for Foldalign compared with CRAST; and several incorrect maps of the target ones to the corresponding shuffled query ones for Foldalign (32) compared with CRAST (0). It also implies alignment with simultaneous folding could detect numerous non-homologous ones rather than sufficient homologs despite the huge computation compared with CRAST.

**Table 2:** performance comparison with Foldalign [9]

3.7 Performance comparison with Foldalign
| Tool/term | TPs/FPs/TNs/FNs | F-meas. | DB[s] | Align.[s] |
| --- | --- | --- | --- | --- |
| CRAFT | 65/156/219/0 | 0.454 | 728.2 | 7.582 |
| Foldalign | 32/1,923/1,923/32 | 0.031 | — | > 3[d] |

We define the "FP" as any target one not mapped to the corresponding query one; the "TN" as any target one not mapped to the corresponding shuffled query one.
Everything except for them is the same as Table 1.

## 4 DISCUSSION

### 4.1 Principal findings

We developed the CRAST algorithm (the time complexity *O*(*n*: *a sum of lengths of target sequences*)) to pairwise-align numerous ncRNAs with consideration of both of sequence/secondary structure identity. Instead of explicit consideration of secondary structure like the Sankoff algorithm, we utilized RNA context (motif probabilities) from the CapR algorithm and fused the score from sequence/secondary structure into the fusion score for the implicit consideration.

We demonstrated it could successfully reduce detections of non-homologous ones keeping detections of homologs as many as other BLAST-like tools with reasonable computation-time in case of lncRNA, in other words, low sequence identity requiring viewpoint of secondary structure identity. This reduction of the false detections leads to improvement of product quality/computation time from subsequent process in genome comparative analysis such as binary classification of ncRNA by RNAz [23, 24] and ncRNA clustering by GraphClust [25].

Surprisingly, Foldalign, a variant of the Sankoff algorithm could detect numerous non-homologous ones rather than sufficient homologs despite the huge computation compared with CRAST in case of lncRNA. It may be because of a fact which only highly similar secondary structure would lead to incomplete capture of homologs as in Fig. 7 and Fig. 9, and even excess capture of non-homologous ones.

To verify it, we set the seed E-value parameter to 0.75; other E-value ones to 1; and contribution ratio of base to score *r* to 0.5 for relaxing/restricting sequence/secondary structure identity and equally taking both of the identities into account in the scoring system. Then we got 31/1,088/1,062/28 as TPs/FPs/TNs/FNs in the same condition as Table 2; the number of the TPs is comparable with the Foldalign result in Table 2. The lower number of the FPs/TNs could be due to adaptive seed requiring rare exact match on sequence.

### 4.2 Method appraisal

We independently modelled a distribution of series of matches and mismatches on sequence/context sequence as a binomial distribution *B*(*n, p*) due to uncertainty of the score distribution/gap. In general, Modelling a distribution of scores as the Gumbel distribution instead of matches is frequently used to handle the uncertainties. But the fusion score depends on not only the match/mismatch score and gap opening/extension cost but also the contribution ratio of base to the fusion score *r* and estimated RNA context sequence controlled by the parameter *w* from the CapR algorithm. We could model it as the distribution by fitting random data for roughly possible combinations of the parameters.

We demonstrated the algorithm between only human and house mouse due to limited availability of sufficient annotation of lncRNA homolog. However, the more evolutional distance on ncRNA between 2 compared species gets diverse, the more homolog detectability of CRAST/other BLAST-like tools/the Sankoff algorithm may increase/decrease. If we got the sufficient availability, we would robustly parameter-tune the algorithm.

We reasoned why the alignment step of CRAST was relatively slow: missing pre-computing of seed candidates (possible adaptive seeds) in the database step and the compute-intensive of the Jensen-Shannon distance. Except for the latter factor, we would implement the former one in CRAST, and let it faster and more accurate.

### 4.3 Scientific implications

We discovered the Sankoff algorithm, a traditional ncRNA alignment could lead to numerous non-homologous ones rather than sufficient homologs and BLAST-like tools work well in case of lncRNA contrary to our expectations. The result and performance of CRAST lead to comparing a large amount of ncRNA pairs in the efficient and accurate fashion. We could adapt the algorithm to ncRNA multiple alignment with consideration of secondary structure such as [26] and ncRNA clustering with the consideration such as [25].

## 5 ACKNOWLEDGEMENTS

The computations were performed on the NIG supercomputer at the ROIS National Institute of Genetics.

We thank Martin C. Frith for telling us about optimization of LAST/BLASTN and the LAST implementation; Anish MS Shrestha for suggesting to represent the ROC curves of the BLAST-like tools for comparison and to parameterize RNA context match probability; Kiyoshi Asai for setting up an opportunity to discuss with them.

## 6 APPENDICES

### 6.1 Default CRAST parameters

**Table 3:** default CRAST parameters from parameter-tuning

| Term | Description | Value |
| --- | --- | --- |
| ms | Maximum span between paired bases for CapR algorithm | 200 |
| ts | Threshold of frequency of adaptive seeds | 3 |
| x1 | Threshold of drop of score when ungapped alignment | 5 |
| x2 | Threshold of drop of score when gapped alignment | 15 |
| cp | Match probability of RNA context | 0.25 |
| b | Contribution ratio of base to score | 0.65 |
| e1 | Threshold of # E-value of seeds with better scores on secondary structure | $6.5 \cdot 10^2$ |
| e2 | Threshold of # E-value of ungapped alignments with better scores on sequence | $5 \cdot 10^{-3}$ |
| e3 | Threshold of # E-value of ungapped alignments with better scores on secondary structure | $9 \cdot 10^2$ |
| e4 | Threshold of # E-value of gapped alignments with better scores on sequence | $5 \cdot 10^{-4}$ |
| e5 | Threshold of # E-value of gapped alignments with better scores on secondary structure | $9 \cdot 10^2$ |
| e6 | Threshold of # E-value of final gapped alignments with better scores on sequence | $5 \cdot 10^{-4}$ |
| e7 | Threshold of # E-value of final gapped alignments with better scores on secondary structure | $6.5 \cdot 10^3$ |
| go | Gap opening penalty | -7 |
| ge | Gap extension penalty | -1 |

E-values $E[M]$ are per target sequence: $E[M] = s(1 - B(M \leq m; n, p))$ where $s$ is a possible number of the seed/alignment occurrence and $M/m$ is a random variable/observed value for a number of matches of bases/secondary structure. We set $ms$ equal to the default one of the CapR algorithm and $go/ge$ equal to the default one of LAST.

### 6.2 Zsh commands for BLAST-like tools/Foldalign

**Listing 1:**
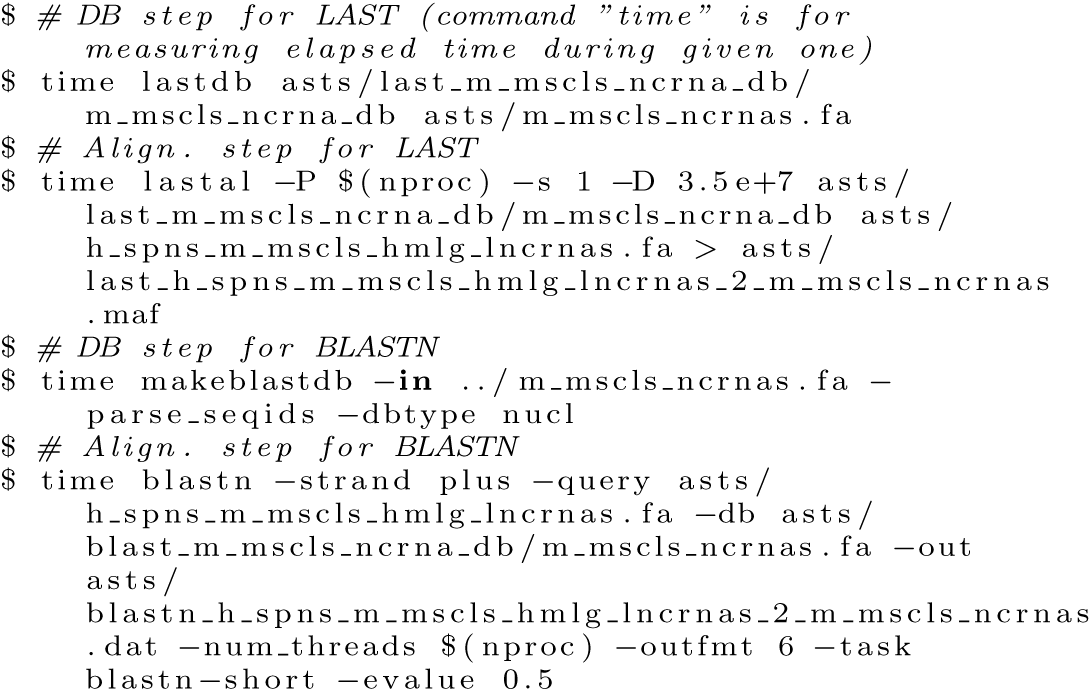
Zsh commands for BLAST-like tools

**Listing 2:**
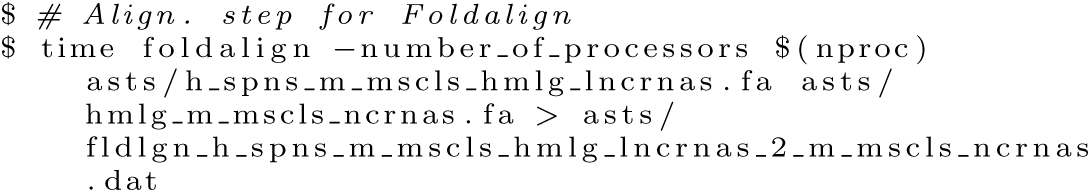
Zsh commands for Foldalign

For the shuffled query sequences, just replace the query file with the one for them, prepending “shfl” to the name of the query file when issuing a command of an alignment step.

